# SARS-CoV-2 viroporin triggers the NLRP3 inflammatory pathway

**DOI:** 10.1101/2020.10.27.357731

**Authors:** Huanzhou Xu, Siddhi A. Chitre, Ibukun A. Akinyemi, Julia C. Loeb, John A. Lednicky, Michael T. McIntosh, Sumita Bhaduri-McIntosh

## Abstract

Cytokine storm resulting from a heightened inflammatory response is a prominent feature of severe COVID-19 disease. This inflammatory response results from assembly/activation of a cell-intrinsic defense platform known as the inflammasome. We report that the SARS-CoV-2 viroporin encoded by ORF3a activates the NLRP3 inflammasome, the most promiscuous of known inflammasomes. ORF3a triggers IL-1β expression via NFκB, thus priming the inflammasome while also activating it via ASC-dependent and -independent modes. ORF3a-mediated inflammasome activation requires efflux of potassium ions and oligomerization between NEK7 and NLRP3. With the selective NLRP3 inhibitor MCC950 able to block ORF3a-mediated inflammasome activation and key ORF3a residues needed for virus release and inflammasome activation conserved in SARS-CoV-2 isolates across continents, ORF3a and NLRP3 present prime targets for intervention.

**Summary:** Development of anti-SARS-CoV-2 therapies is aimed predominantly at blocking infection or halting virus replication. Yet, the inflammatory response is a significant contributor towards disease, especially in those severely affected. In a pared-down system, we investigate the influence of ORF3a, an essential SARS-CoV-2 protein, on the inflammatory machinery and find that it activates NLRP3, the most prominent inflammasome by causing potassium loss across the cell membrane. We also define key amino acid residues on ORF3a needed to activate the inflammatory response, and likely to facilitate virus release, and find that they are conserved in virus isolates across continents. These findings reveal ORF3a and NLRP3 to be attractive targets for therapy.

## Main Text

Worldwide reports of COVID-19 indicate that effective management of severely ill individuals will require both antiviral and anti-inflammatory strategies. Indeed, during the second week of illness, those with severe disease experience cytokine storms indicating a massive inflammatory surge ^1,2^. This inflammatory response, composed of IL-1β and other cytokines, results from assembly/activation of a multiprotein host machinery known as the inflammasome in both immune and non-immune cells such as airway epithelial cells – the most prominent is the NLRP3 (NOD-, LRR- and pyrin domain-containing protein 3)-inflammasome – and several lines of evidence tie activation of the NLRP3-inflammasome to severe SARS-CoV-2 pathology, including i) individuals with comorbidities such as diabetes, atherosclerosis, and obesity (all pro-inflammatory conditions marked by NLRP3 activation) ^3–8^ are at greater risk for severe disease ^9–11^, ii) viroporins expressed by the closely-related SARS-CoV activate the NLRP3 inflammasome ^12^, and iii) bats, the asymptomatic reservoir of CoVs that are highly pathogenic in humans, are naturally defective in activating the NLRP3-inflammasome ^13^. Although cellular ACE2 engagement by SARS-CoV-2 spike protein can cause expression of pro-inflammatory genes ^14^, whether CoV-2 activates the inflammasome remains unexplored.

Given the central role of inflammation in severe COVID-19 and the high level of conservation of the viroporin ORF3a across CoV genomes, we investigated the influence of the SARS-CoV-2 ORF3a on the NLRP3 inflammasome. Viroporins are virus-encoded proteins that are considered virulence factors. Though typically not essential for virus replication, some of these small hydrophobic proteins can form pores that facilitate ion transport across cell membranes, and by so doing, ensure virus release with the potential for coincident inflammasome activation ^15,16^. A component of the innate immune system, the inflammasome assembles and responds to invading organisms, thus forming the first line of defense against infections ^17^. Our experiments show that the CoV-2 ORF3a protein primes and activates the inflammasome via efflux of potassium ions and the kinase NEK7. Its ability to activate caspase 1, the central mediator of proinflammatory responses, depends on NLRP3 since a selective inhibitor of NLRP3 blocks this pathway in infected cells. Importantly, we find that although the CoV-2 ORF3a protein has diverged somewhat from its homologs in other CoVs, some of these newly divergent residues are essential for activating the NLRP3 inflammasome and are perfectly conserved in virus isolates across continents.

### SARS-CoV-2 viroporin ORF3a primes and activates the inflammasome, prompting cell death

With lung as the predominant site of pathology along with established tropism for kidney and other organs ^18^, we introduced ORF3a into lung origin A549 cells and for comparison, kidney origin HEK-293T cells, cell types that readily support SARS-CoV-2 infection ^19^, and found induction of pro-IL-1β in both cell types, consistent with priming of the inflammasome. Compared to empty vector-exposed cells, ORF3a also increased the levels of cleaved, i.e. the active form of the pro-inflammatory caspase, caspase 1, as well as the cleaved form of the caspase 1 substrate, pro-IL-1β, indicating activation of the inflammasome, again in both cell types (Fig.1A). Priming by ORF3a resulted from NFκB-mediated expression of *IL-1β* message (Fig.1B) as indicated by increased IκBα phosphorylation and enrichment of NFκB p65 at the *IL-1β* promoter in ORF3a-exposed cells (Figs.1C-E). ORF3a also caused cleavage/activation of Gasdermin D, the pyroptosis-inducing caspase 1-substrate, indicated by an increase in the N-terminal fragment of Gasdermin D (Fig.1F). This was accompanied by ORF3a-mediated increased cleavage/activation of caspase 3 and cell death, likely secondary to both pyroptosis and apoptosis (Figs.1G and H). Thus, ORF3a primes the inflammasome by triggering NFκB-mediated expression of pro-IL-1β while also activating the inflammasome to cleave pro-caspase-1, pro-IL-1β, and the pore-forming Gasdermin D, inducing cell death.

**Figure 1.**
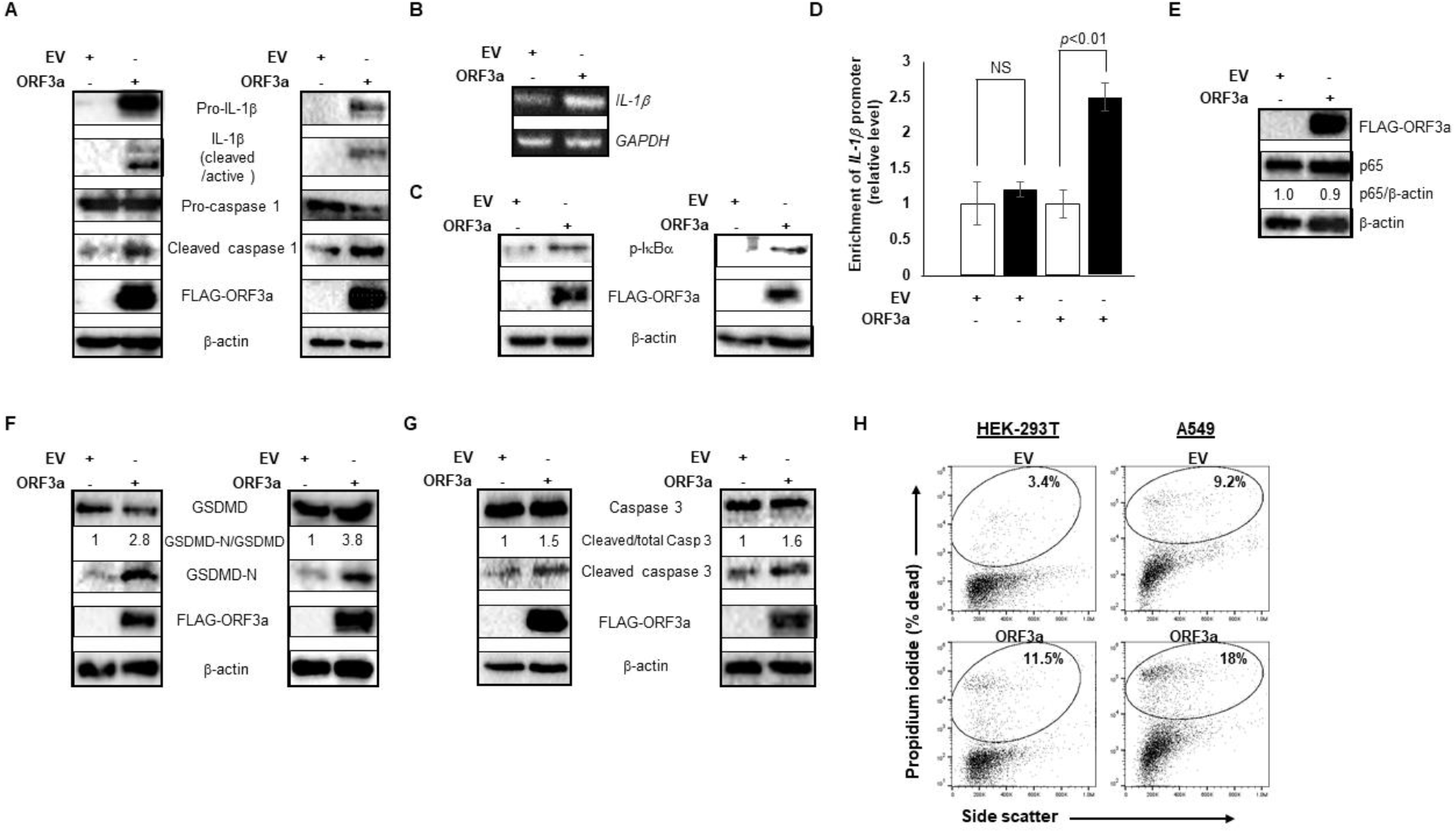
ORF3a primes and activates the inflammasome, causing cell death. HEK-293T cells (A, C, F, G; left panels) or A549 cells (A, C, F, G; right panels) were transfected with FLAG-tagged ORF3a or empty vector (EV) and harvested after 24 hours for immunoblotting with indicated antibodies. (B, D, E) ORF3a-transfected A549 cells were analyzed at 24 hours by reverse transcriptase-quantitative PCR for *IL-1β* mRNA abundance (B), ChIP-PCR to quantify relative enrichment of NFkB p65 at the *IL-1β* promoter using anti-p65 antibodies (black bar) or control IgG (white bar) (D), or immunoblotting as indicated (E). Unfixed cells harvested 24 hours after transfection with EV or ORF3a were stained with propidium iodide followed by flow cytometry to enumerate percent dead cells in H. Error bars in D represent SEM. All experiments were performed three times.

### ORF3a activates the NEK7-NLRP3 inflammasome via ASC-dependent and independent modes

In probing the mechanism of ORF3a-mediated activation of the inflammasome, we found that it enhanced NLRP3 protein levels, and knockdown of NLRP3 curbed ORF3a-directed caspase 1 cleavage (Figs.2A-B), indicating priming and activation of the NLRP3 inflammasome by ORF3a. Further, MCC950, a selective small molecule inhibitor that binds to the NACHT domain of NLRP3 and curtails its activation by blocking ATP hydrolysis ^20^, also blocks ORF3a-mediated activation of the inflammasome in low micromolar concentrations (Fig.2C). Moreover, with the NIMA-related kinase NEK7 recently linked to NLRP3 activation ^16^, we also depleted NEK7 and found that ORF3a was impaired in its ability to cause cleavage of caspase 1, i.e. unable to activate the inflammasome (Fig.2D). The NLRP3 inflammasome is activated by a variety of cell-extrinsic and -intrinsic stimuli that trigger the assembly of the inflammasome machinery wherein NLRP3 oligomerizes with the adaptor protein ASC (Apoptosis-associated speck-like protein containing a CARD) leading to recruitment of pro-caspase 1 which is then activated by proximity-induced intermolecular cleavage. Given ORF3a-mediated inflammasome activation in HEK-293T cells that lack ASC (Fig.2E), we asked if ORF3a activated the inflammasome solely in an ASC-independent manner. We found that ORF3a’s ability to activate pro-caspase 1 was substantially impaired upon depletion of ASC in A549 cells (Fig.2F), supporting the idea that ORF3a activates the inflammasome in both ASC-dependent and -independent ways. To assess if ORF3a also mediates activation of other prominent inflammasomes including NLRP1 and NLRC4, we depleted each of these molecules but were unable to block cleavage of pro-caspase 1 (Fig.2G), indicating that ORF3a predominantly activates the NLRP3 inflammasome.

**Figure 2.**
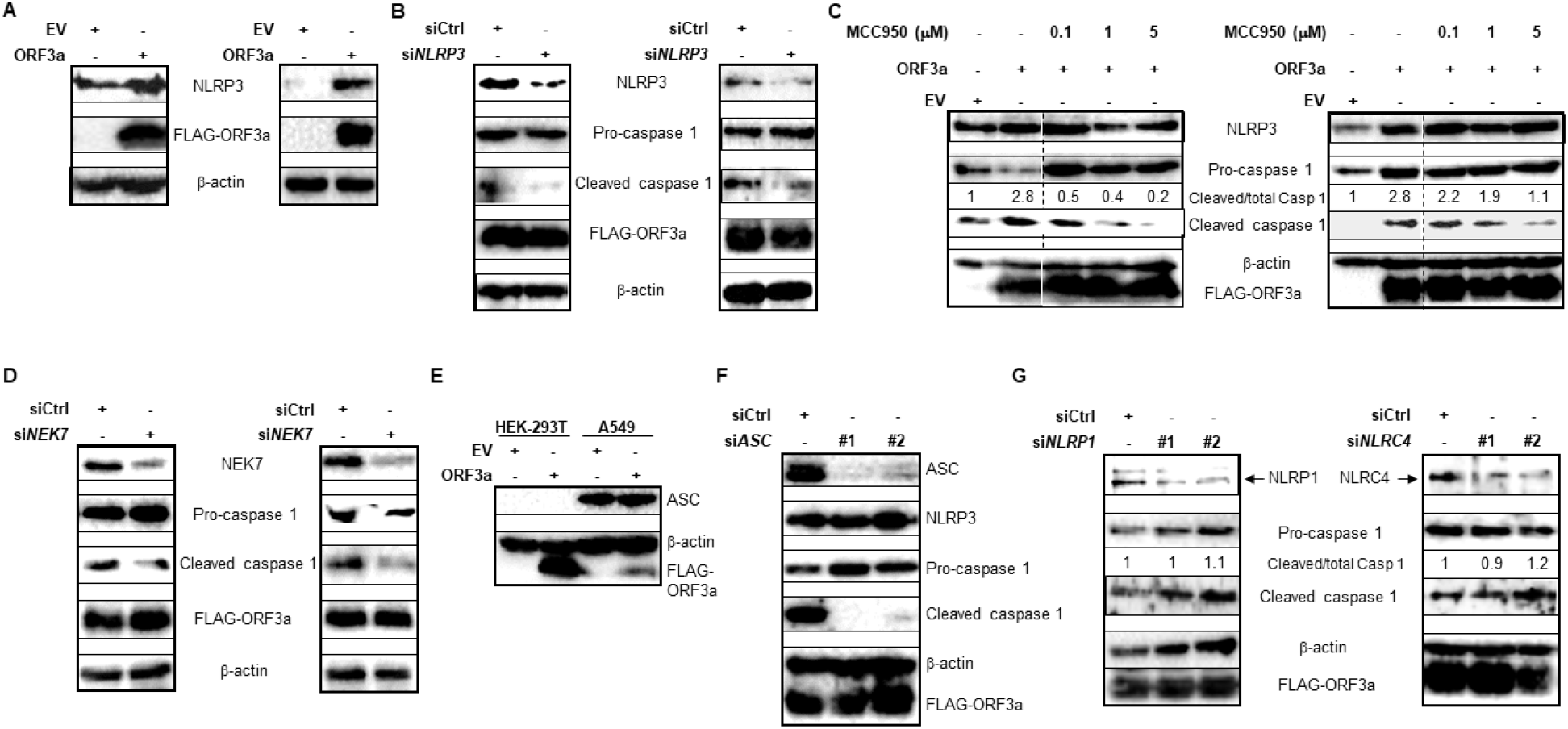
ORF3a activates the NEK7-NLRP3 inflammasome via ASC-dependent and independent modes. (A) Cell lysates of FLAG-ORF3a- or EV-transfected HEK-293T (left) or A549 cells (right) were immunoblotted with indicated antibodies. (B, D) HEK-293T (left) or A549 cells (right) were co-transfected with FLAG-ORF3a and control siRNA (B, D), *NLRP3* siRNA (B), or *NEK7* siRNA (D) for 24 hours prior to immunoblotting with indicated antibodies. (C) HEK-293T (left) or A549 cells (right) were transfected with EV or FLAG-ORF3a and exposed to MCC950 for 24 hours prior to immunoblotting. (E) Cell lysates were immunoblotted with indicated antibodies. (F, G) A549 cells were co-transfected with FLAG-ORF3a and control siRNA, *ASC* siRNA (F), *NLRP1* siRNA (G; left), or *NLRC4* siRNA (G, right) for 24 hours prior to immunoblotting with indicated antibodies. Experiments were performed at least thrice.

### ORF3a triggers NLRP3 inflammasome assembly via K^+^ efflux

With NEK7 a key mediator of NLRP3 activation downstream of potassium efflux, and efflux of potassium ions a central mechanism of NLRP3 activation, particularly by ion channel-inducing viroporins ^15,16,21^, we investigated the effect of blocking potassium efflux by raising the extracellular concentration of K^+^ and found that ORF3a-mediated caspase 1 cleavage was abrogated (Fig.3A). To identify the type of K^+^ channel formed by ORF3a, we employed known pharmacologic inhibitors including quinine, barium, iberiotoxin, and tetraethylammonium to block two-pore domain K^+^ channels, inward-rectifier K^+^ channels, large conductance calcium-activated K^+^ channels, and voltage gated K^+^ channels, respectively ^22^. Mimicking the ability of barium to block the release of SARS-CoV virions ^23^ and supporting the finding in Fig.3A, barium was able to curb CoV-2 ORF3a-mediated activation of caspase 1, indicating that ORF3a forms inward-rectifier K^+^ channels in the cell membrane (Fig.3B). Restricting K^+^ efflux also impaired ORF3a’s ability to trigger assembly of both ASC-independent and -dependent NLRP3 inflammasomes (Figs.3C and D, respectively). Notably, not only did SARS-CoV-2 activate the inflammasome upon infection of A549 and HEK-293T cells, but this activation was dampened by MCC950 and blocking K^+^ efflux (Figs.3E and F), asserting the importance of ion channels and NLRP3 in triggering the inflammatory response in CoV-2 infected cells.

**Figure 3.**
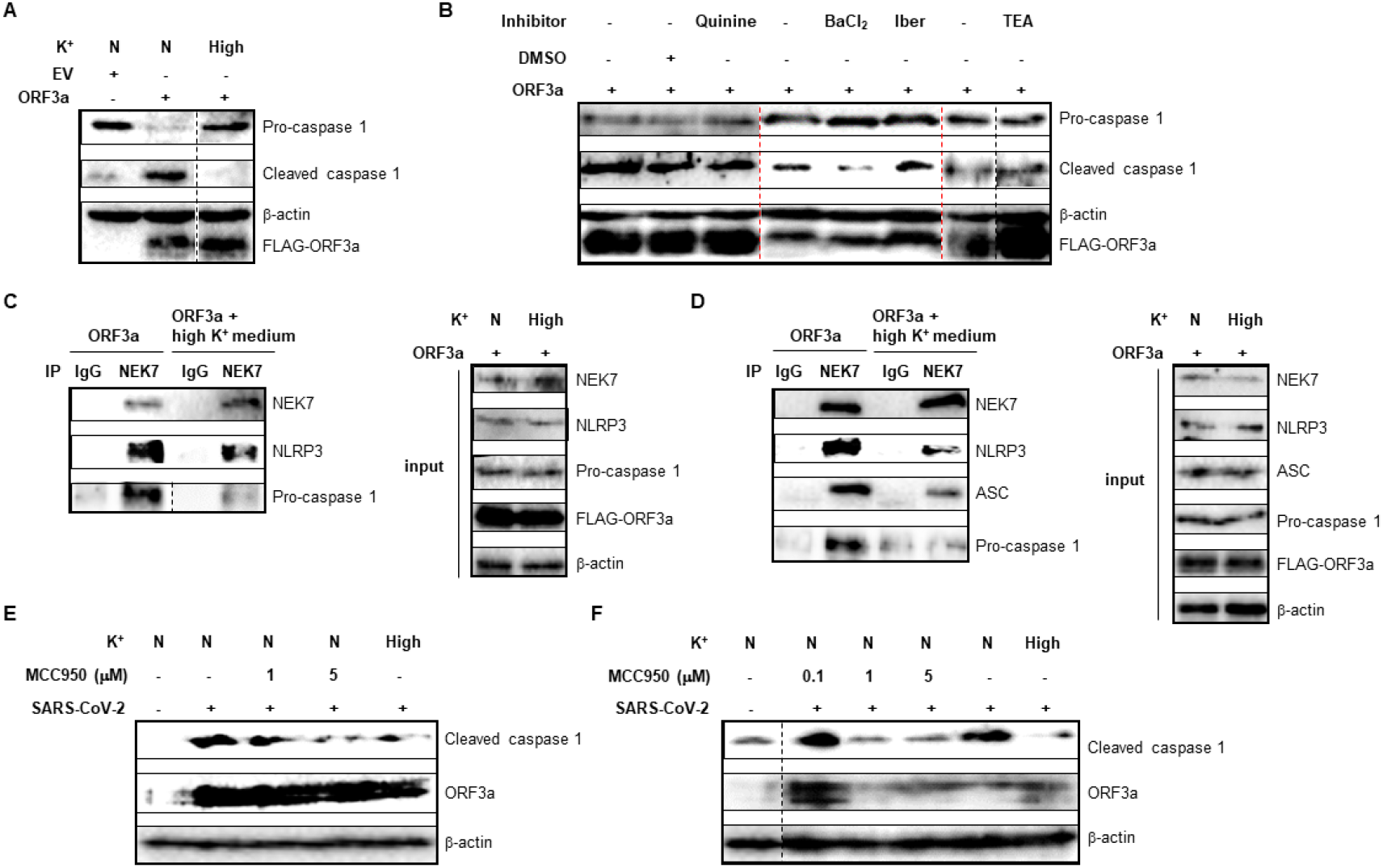
ORF3a-mediated activation of NLRP3 inflammasome requires K^+^ efflux. (A) FLAG-ORF3a plasmid or EV were introduced into A549 cells. After 20 hours, cells were left in normal medium (N) or exposed to medium with high K^+^ (50mM; to block K^+^ efflux; High). Cells were harvested 4 hours later, and extracts immunoblotted with indicated antibodies. (B) A549 cells transfected with FLAG-ORF3a were exposed to indicated potassium channel inhibitors quinine, barium (BaCl2), iberiotoxin (Iber), and tetraethylammonium (TEA) for 24 hours prior to immunoblotting with different antibodies. (C and D) FLAG-ORF3a plasmid was introduced into HEK-293T (C) and A549 (D) cells. After 20 hours, cells were left in normal medium (N) or exposed to medium with high K^+^ (High). Cells were harvested 4 hours later, and extracts immunoblotted (Input) or immunoprecipitated with control IgG or anti-NEK7 antibody followed by immunoblotting with indicated antibodies. Input represents 5% of sample. (E and F) HEK293T (E) and A549 (F) cells were infected with SARS-CoV-2 in the presence of MCC950 or high K^+^containing medium (High; for the last 20 hours of culture) and harvested after 24 hours for immunoblotting with indicated antibodies. Experiments were performed twice.

### Key residues in ORF3a important for activating the inflammasome are well conserved

Alignment of ORF3a sequences from SARS-CoV-2 isolates from Asia, Europe, Middle-East, Russia, and North and South America between December 2019 and June 2020 as well as other bat CoVs and SARS-CoV revealed the conservation of two out of three key cysteine residues (residues 127, 130, and 133), shown to be essential for K^+^ channel formation by SARS-CoV ^21^ (Fig.4A). The exception, cysteine 127, was replaced by leucine in all CoV-2 isolates. We also observed a similar switch from cysteine to valine at position 121 and a switch from asparagine to cysteine at position 153 in all CoV-2 isolates. Introducing single point mutations at positions 127, 130, and 133 of CoV-2 ORF3a impaired its ability to activate the inflammasome, supporting the need for not only the two conserved cysteines at positions 130 and 133 but also that of the newly acquired leucine at position 127 of CoV-2 ORF3a (Fig.4B). Similarly, mutating the residues at positions 121 and 153, both newly acquired in CoV-2 though conserved in all isolates, resulted in a dampened response by the inflammasome (Fig.4B). Thus, SARS-CoV-2 ORF3a has retained some of the key residues needed for virus release and inflammasome activation but it has acquired additional changes that support a functionally consequential divergence from earlier CoVs. Nonetheless, this domain bearing the abovementioned residues that is essential for forming ion channels for virus release has remained remarkably well conserved throughout the pandemic, thereby maintaining its ability to activate the inflammasome.

**Figure 4.**
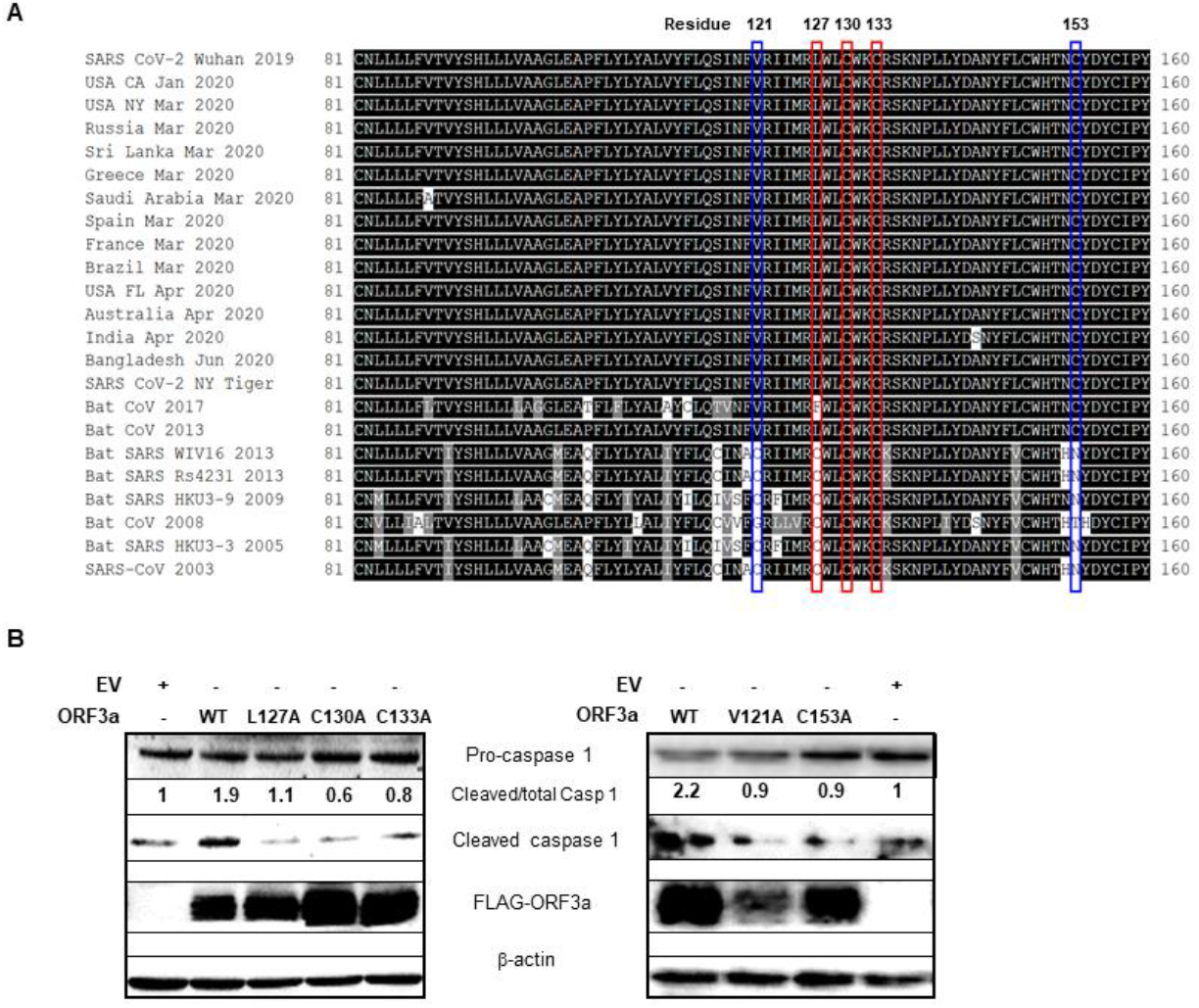
ORF3a residues required for inflammasome activation are conserved in SARS-CoV-2 isolates across continents. (A) ORF3a/ORF3 viroporin from SARS-like betacoronaviruses including temporally and geographically distinct isolates from the COVID-19 pandemic and diverse species isolates dating back to the original SARS pandemic of 2003 were aligned in CLUSTAL Omega using EMBL-EBI Server Tools (https://www.ebi.ac.uk/Tools/services/web_clustalo/toolform.ebi). Selected isolates displaying the most diversity are shown from positions 81 to 160 of ORF3a/ORF3. Conserved cysteine residues previously identified in SARS-CoV as critical to K^+^ ion channel formation are outlined in red. Newly divergent residues (121 and 153) conserved across SARS-CoV-2 isolates are outlined in blue. The multiple sequence alignment was shaded in BoxShade hosted by ExPASy (https://embnet.vital-it.ch/software/BOX_form.html). (B) A549 cells were transfected with EV, wild-type FLAG-ORF3a (WT), or FLAG-ORF3a mutants. Cells were harvested 24 hours later and immunoblotted with indicated antibodies. Experiments were performed at least twice.

## Discussion

In summary, an essential viroporin required for release of SARS-CoV-2 from infected cells is also able to prime and activate the NLRP3 inflammasome, the machinery responsible for much of the inflammatory pathology in severely ill patients. ORF3a’s indispensability to the virus’s life cycle makes it an important therapeutic candidate. Moreover, while different from its homologs in other CoVs, the high conservation of the newly divergent SARS-CoV-2 ORF3a across isolates from several continents combined with our observation that multiple single point mutations reduce its ability to activate the inflammasome, argues against rapid emergence of resistance phenotypes. Thus, targeting ORF3a has the dual potential of blocking virus spread and inflammation.

SARS-CoV-2 is not only linked to severe and fatal outcomes in adults with underlying comorbidities associated with pre-existing inflammation, but it also causes severe disease in children in the form of Multisystem Inflammatory Syndrome in Children (MIS-C) as well as in adults as MIS-A ^24,25^. Dampening the inflammatory response in such patients is therefore an attractive strategy – a strategy that has shown promise in a small group of patients treated with Anakinra, a recombinant IL-1R antagonist ^26^. Notably, for several inflammatory diseases, there is keen interest within the pharmaceutical industry in therapeutically targeting the inflammatory pathway at a further upstream point, namely NLRP3 itself. MCC950 is a prototype of this approach with several other related compounds undergoing preclinical, phase I, and phase II trials (https://cen.acs.org/pharmaceuticals/drug-discovery/Could-an-NLRP3-inhibitor-be-the-one-drug-to-conquer-common-diseases/98/i7). Along the same lines, Gasdermin D, also activated by ORF3a, presents yet another therapeutic target as it may potentiate virus release by killing cells in addition to causing inflammation. Additionally, restraining the NLRP3 inflammasome may secondarily stifle virus replication itself as we recently demonstrated for a DNA tumor virus ^27^.

Aside from ORF3a, other viroporins such as ORF-E and ORF8 may also contribute to the inflammatory response by similar or related mechanisms. Activation of the NLRP3 inflammasome also bears mention in broader contexts. In particular, two recent reports have found that a fraction of severely ill COVID-19 patients display defective type I interferon immunity ^28,29^. It is likely that severe disease in these individuals also stemmed from unchecked pro-inflammatory responses since type I interferon can counteract the NLRP3 inflammasome in a number of ways ^30^. Similarly, for those who have metabolic disturbances such as hypokalemia that often results from antihypertensive medications, ORF3a may have a lower threshold for activating the inflammasome due to a higher K^+^ gradient across the infected cell.

## Author contributions

Conceptualization: H.X., S.B.-M.; Methodology: H.X., M.T.M., S.B.-M.; Validation: H.X., S.B.-M.; Formal analysis: H.X., S.B.-M.; Investigation: H.X., S.A.C., I.A.A., J.G., J.L., M.T.M.; Writing – original draft preparation: H.X., S.B.-M.; Writing – review and editing: M.T.M., S.B.-M.; Visualization: H.X., S.B.-M., M.T.M.; Supervision: S.B.-M.; Project administration: S.B.-M., Funding acquisition: S.B.-M.

## Data and materials availability

All data is available in the main text or the supplementary materials.

## Methods

### Cell lines and infection

Human embryonic kidney-293T (HEK-293T) cells were maintained in DMEM (Thermo Fisher Scientific, Cat. 11965118) containing 10% fetal bovine serum (GEMINI, Cat. 900108) and 1% penicillin/streptomycin (Gibco, Cat. 15140122). A549 cells were maintained in Ham’s F-12 Nutrient Mix (Thermo Fisher Scientific, Cat. 11765054) containing 10% fetal bovine serum and 1% penicillin/streptomycin. Both cell lines were cultured in the presence of 5% CO2 at 37 °C. Cells were infected in a BSL-3 lab with the UF-1 strain of SARS-CoV-2 at MOI of 4 in media containing 3% low IgG FBS (Fisher Scientific, Cat. SH30070.03).

### Plasmids, siRNAs, and transfection

The *ORF3a* gene without stop codon (nt 25,382-26,206, GenBank accession no. MT295464.1) was PCR amplified with forward primer (5’CGCGGATCCATGGATTTGTTTATGAGAATCTT3’) and reverse primer (5’ AAGGAAAAAAGCGGCCGCCAAAGGCACGCTAGTAGTC3’) by using Phusion High-Fidelity DNA Polymerase (New England Biolabs, M0530L) according to the manufacturer’s protocol and inserted into pcDNA5.1/FRT/TO vector (a kind gift from professor Torben Heick Jensen, Denmark) with a C-terminal 3×FLAG tag to generate FLAG-tagged ORF3a plasmid. Flag-tagged ORF3a mutants (V121A, L127A, C130A, C133A, C153A) were constructed by overlap extension PCR with the following primer pairs:

5’GAGTATAAACTTTGCAAGAATAATAATGAG3’ (forward) and

5’ CTCATTATTATTCTTGCAAAGTTTATACTC3’ (reverse),

5’ATAATGAGGGCTTGGCTTTG3’ (forward) and

5’ CAAAGCCAAGCCCTCATTAT3’ (reverse),

5’GCTTTGGCTTGCCTGGAAATGC3’ (forward) and

5’ GCATTTCCAGGCAAGCCAAAGC3’ (reverse),

5’TGCTGGAAAGCCCGTTCCAAA3’ (forward) and

5’ TTTGGAACGGGCTTTCCAGCA3’ (reverse),

5’GCATACTAATGCTTACGACTATTG3’ (forward) and

5’CAATAGTCGTAAGCATTAGTATGC3’ (reverse), respectively.

HEK-293T and A549 cells were transfected with LipoJet™ In Vitro Transfection Kit (SignaGen Laboratories, SL100468) according to the manufacturer’s protocol.

HEK-293T and A549 cells were transfected with 200 pmoles of siRNA. siRNAs included *NLRP3* (Ambion, Cat. s41554), *NEK7* (Ambion, Cat. 103794), *ASC* (Ambion, Cat. 44232 and 289672), *NLRP1* (#1, Ambion, Cat. S22520; #2, Ambion, Cat. 239345), *NLRC4* ((#1, Ambion, Cat. S33828; #2, Ambion, Cat. 105219), and control (Dharmacon, Cat. D001810-01-20).

### SARS-CoV-2

A passage two stock of *Severe acute respiratory syndrome coronavirus 2* isolate SARS-CoV-2/human/USA/UF-1/2020 (GenBank MT295464) was used for virus-infection studies. The virus was the first isolate from a patient at the University of Florida Health Shands Hospital (J. Lednicky, unpublished) and has about 99% nt identity with SARS-CoV-2 reference strain Wuhan-Hu-1 (GenBank NC_045512.2) and 100% identity with the genomes of SARS-CoV-2 detected in California, USA. The genome of SARS-CoV-2 UF-1 encodes an aspartic acid residue at amino acid 614 of the spike protein. This virus was isolated and then propagated (one passage) in VeroE6 cells prior to sequence analyses and use in this work, and has no INDELs in its genome. All work with this virus was performed in a BSL-3 laboratory by an analyst using a full-head powered-air purifying respirator and appropriate personal protective equipment, including gloves and a chemically impervious Tyvek gown.

### Chemical treatment of cell lines

HEK-293T and A549 cells were transfected with plasmids. After 2h, different chemical reagents were added to medium. Chemical reagents included NLRP3 inhibitor MCC950 (0.1–5 μM) (Sigma Aldrich, Cat. 538120), Quinine (10 μM) (Sigma Aldrich, Cat. 145904), Barium chloride (2 mM) (Sigma Aldrich, Cat. 342920), Tetraethylammonium chloride (5 mM) (Tocris Bioscience, Cat. 306850), and Iberiotoxin (1.0 μM) (Tocris Bioscience, Cat. 1086100U). All chemicals were dissolved with DMSO or sterile water.

### Reverse transcription PCR (RT-PCR)

RT-PCR was performed as previously described ^1^. Briefly, 1 μg of total RNA was used as template for complementary DNA synthesis using MuLV reverse transcriptase (New England Biolabs, Cat. M0253L) according to the manufacture’s protocol. OneTaq DNA Polymerase (New England Biolabs, Cat. M0480S) was used to amplify DNA fragment using manufacture’s protocol. RT-PCR primers were as following:

forward primer 5’ACCATCTTCCAGGAGCGAGA3’ and

reverse primer 5’GGCCATCCACAGTCTTCTGG 3’ for *GAPDH* mRNA,

forward primer 5’TCAGCCAATCTTCATTGCTC3’ and

reverse primer 5’GCCATCAGCTTCAAAGAACA3’ for *IL-1β* pre-mRNA ^2^

### Immunoblotting and antibodies

Immunoblotting was performed as previously described ^3^. Briefly, total cell lysates were electrophoresed on 10% or 12% SDS-polyacrylamide gels and transferred onto nitrocellulose membranes and immunoassayed with indicated antibodies. The following antibodies were used: rabbit anti-Caspase-1 antibody (Thermo Scientific, Cat. PA587536), rabbit anti-cleaved Caspase-1 antibody (Thermo Scientific, Cat. PA538099), mouse anti-IL-1β (Cell Signaling Technology, Cat. 12242s), mouse anti-Flag M2 antibody (Sigma-Aldrich, Cat. F1804), mouse anti-β-actin antibody (Sigma-Aldrich, Cat. A5441), rabbit anti-phospho-IκBα (Ser32) antibody (Cell Signaling Technology, Cat. 2859s), rabbit anti-IκBα antibody (Cell Signaling Technology, Cat. 9242s), rabbit anti-NF-κB p65 antibody (Cell Signaling Technology, Cat. 8242s), rabbit anti-Caspase 3 antibody (GeneTex, Cat. GTX110543), rabbit anti-Gasdermin D (L60) antibody (Cell Signaling Technology, Cat. 93709s), rabbit anti-cleaved-Gasdermin D (Asp275) antibody (Cell Signaling Technology, Cat. 36425s), rabbit anti-NLRP3 antibody (Invitrogen, Cat. PA5-21745), rabbit anti-NEK7 antibody (Cell Signaling Technology, Cat. 3057s), rabbit anti-ASC antibody (Cell Signaling Technology, Cat. 13833s), rabbit anti-NLRP1 antibody (Novus Biologicals, Cat. NB100-56147SS), rabbit anti-NLRC4 antibody (Novus Biologicals, Cat. NB100-56142SS), rabbit anti-SARS-CoV-2 ORF3a antibody (FabGennix, Cat. SARS-COV2-ORF3A-101AP), HRP-conjugated goat anti-mouse IgG(H+L) (Thermo Scientific, Cat. 626520) and HRP conjugated goat anti-rabbit IgG(H+L) (Thermo Scientific, Cat. 31460), and HRP conjugated goat anti-rabbit IgG (light chain) (Novus, Cat. NBP2-75935).

### Flow cytometry

Flow cytometry was performed as previous described ^4^. Briefly, HEK-293T and A549 cells were treated with trypsin for 3 min and collected by centrifugation at 350g for 3 min. Cell pellets were washed twice with FACS buffer (1X PBS with 2% FBS) and resuspended in 200 μl of RNase-containing FACS buffer. 20 μl of propidium iodide (10 μg/ml) (Sigma-Aldrich, Cat. P4864) was added to each sample and subjected to flow cytometry immediately to assay cell death.

### Chromatin immunoprecipitation-quantitative PCR (ChIP-qPCR)

ChIP was performed as described previously ^5^. Briefly, A549 cells were transfected with FLAG-ORF3a or empty vector as control. Twenty-four hours later, cells (7.5×10^5^ cells for each ChIP) were crosslinked with 1% formaldehyde for 20 min and quenched with 0.125 M glycine. Cells were lysed in 500 μl of nuclear extraction buffer A (Cell Signaling Technology, Cat. 7006) on ice for 15 min and washed once with 500 μl of nuclear extraction buffer B (Cell Signaling Technology, Cat. 7007), and then treated with 0.5 μL of micrococcal nuclease (Cell Signaling Technology, Cat. 10011) for 20 min at 37 °C. Nuclei were resuspended in 1×ChIP buffer (Cell Signaling Technology, Cat. 7008) and sonicated at 8W with 10-s on and 20-s off pulses on ice for two cycles to break nuclear membranes. After removing debris, 2% of each sample was set aside as input and the rest (98%) of the sample was incubated with 3 μg of antibody (or 3 μg of IgG as control) and 30 μl of protein G magnetic beads (Cell Signaling Technology, Cat. 7008) at 4 °C overnight. Beads were washed three times with low salt ChIP buffer and once with high salt ChIP buffer. The protein-DNA complex were eluted with 1×Elution buffer (Cell Signaling Technology, Cat. 10009). DNA was extracted with DNA purification columns (Cell Signaling Technology, Cat. 10010) and subjected to qPCR analysis. The following primers were used for amplifying the *IL-1β* promoter:

forward primer 5’AGGAGTAGCAAACTATGACAC3’ and

reverse primer 5’ACGTGGGAAAATCCAGTATTT3’ ^6^.

### Co-Immunoprecipitation (Co-IP)

Co-IP was performed as described previously ^7^. Cells were lysed in ice-cold IP Lysis Buffer (Thermo Scientific, Cat. 87787) in the presence of 1 ×protease inhibitor cocktail (Cell Signaling, #7012) for 15min followed by centrifugation (14,000 rpm) at 4 °C for 5 min. Of pre-cleared cell lysates, 5% was set aside as input. The rest was incubated with 3.0 μg of rabbit anti-NEK7 antibody (Bethyl Laboratories, Cat. A302-684A) or the same amount of control IgG (R&D, Cat. AB-105-C) together with 40 μl of Dynabeads Protein G (Thermo Scientific, Cat. 10003D) at 4 °C overnight. Beads were washed three times with IP lysis buffer and subjected to immunoblotting.

### Statistical analysis

Unpaired Student’s t test was used to calculate *p* values by comparing the means of two groups.

